# Single cell characterization of a synthetic bacterial clock with a hybrid feedback loop containing dCas9-sgRNA

**DOI:** 10.1101/2020.07.16.206722

**Authors:** John Henningsen, Matthaeus Schwarz-Schilling, Andreas Leibl, Joaquin A. M. Guttierez, Sandra Sagredo, Friedrich C. Simmel

## Abstract

Genetic networks that generate oscillations in gene expression activity are found in a wide range of organisms throughout all kingdoms of life. Oscillatory dynamics facilitates the temporal orchestration of metabolic and growth processes inside cells and organisms, as well as the synchronization of such processes with periodically occurring changes in the environment. Synthetic oscillator gene circuits such as the ‘repressilator’ can perform similar functions in bacteria. Until recently, such circuits were mainly based on a relatively small set of well-characterized transcriptional repressors and activators. A promising, sequence-programmable alternative for gene regulation is given by CRISPR interference (CRISPRi), which enables transcriptional repression of nearly arbitrary gene targets directed by short guide RNA molecules. In order to demonstrate the use of CRISPRi in the context of dynamic gene circuits, we here replaced one of the nodes of a repressilator circuit by the RNA-guided dCas9 protein. Using single cell experiments in microfluidic reactors we show that this system displays robust relaxation oscillations over multiple periods and over the time course of several days. Through statistical analysis of the single cell data, the potential for the circuit to act as a synthetic pacemaker for cellular processes is evaluated. The use of CRISPRi in the context of an oscillator circuit is found to have profound effects on its dynamics. Specifically, irreversible binding of dCas9-sgRNA appears to prolong the period of the oscillator. Further, we demonstrate that the oscillator affects cellular growth, leading to variations in growth rate with the oscillator’s frequency.

---

Over the past two decades, researchers in synthetic biology have created a wide variety of synthetic gene circuits with designed dynamic, sensory or computational functions [1, 2]. Among these, synthetic genetic oscillators are particularly interesting, as they do not only require control over gene expression levels, but also over expression dynamics. In their seminal work, Elowitz and Leibler [3] created a genetic oscillator termed ‘repressilator’, whose negative feedback loop was composed of three genetic repressors (LacI, TetR, and *λ*-cI), which cyclically repressed each other’s expression. Later, a variety of other oscillator architectures were demonstrated, e.g., two-node oscillators containing an activating and a repressing link [4, 5], a mammalian oscillator with negative feedback based on antisense RNA transcription [6], and even a bacterial five-node oscillator containing five cyclically arranged repressor modules [7]. More recently, the dynamical properties of the original repressilator were analyzed in depth, leading to an improved circuit design, which resulted in long-term synchronized oscillations with reduced noise [8].

Oscillatory biochemical reactions naturally occur in a variety of biological contexts [9–11], e.g., in the control of bacterial cell division [12], the coordination of the eukaryotic cell cycle [13, 14], or in circadian rhythms [15]. Similarly, synthetic oscillator circuits could serve as pacemakers or timers for engineered processes. In fact, several potential applications of synthetic oscillators have been recently demonstrated, for instance, to synchronize lysis of therapeutic bacteria for *in vivo* delivery applications [16], or to probe bacterial growth dynamics in the gut [17].

Generally speaking, oscillatory dynamics are generated by non-linear negative feedback loops with a delay [11], where the delay may be achieved, e.g., by distributing the feedback over several intermediate nodes. Non-linearities are required to destabilize a system during its approach towards steady state, and shorter feedback loops have been shown to require stronger non-linearities than longer ones. Apart from non-linearities generated by cooperative binding, also thresholding by molecular titration can be employed, typically resulting in relaxation oscillations [8].

As an exciting alternative to gene regulation via transcription factors, recently the CRISPR interference (CRISPRi) technique has been established, which allows sequence-programmable gene silencing based on the catalytically inactive mutant dCas9 of the CRISPR associated protein Cas9 [18, 19]. Until now, CRISPRi has only rarely been used in the context of dynamical gene circuits, however. Notable exceptions are two recent examples of oscillating networks, in which three single guide RNAs (sgRNAs) cyclically repressed each other’s transcription in the presence of dCas9 [20, 21] or dCas12a [21]. In the present work, we took a hybrid approach – which we termed ‘Rock Paper CRISPR’ (RPC) oscillator-, in which only one of the transcriptional repressor links of a repressilator network was replaced by CRISPRi. Even though CRISPRi is not expected to provide highly non-linear repression, we demonstrate, both experimentally and via computational modeling, that our system is still capable of sustained temporal oscillations.

Long-term monitoring of oscillations of a large number of individual bacteria using a microfluidic ‘mother machine’ [22] allowed us to analyze the dynamic variability and stability of the CRISPRi-modified oscillator at the single cell level.

While direct integration of a CRISPRi process into the repressilator opens up the possibility to control arbitrary genomic functions of the bacteria [23] with a synthetic genetic clock, it may also generate challenges due to the large load put on the cells by CRISPRi. We therefore also investigated aspects such as de-synchronization and phase stability of the oscillator, and its relation to bacterial growth. Using an information-theoretic approach, we analyze its potential as a timekeeper or as a controller for temporal gene expression programs.

## RESULTS

### Design of the gene circuit

Starting with the original repressilator circuit, which contains the TetR, *λ*-phage cI and LacI transcriptional repressors in a negative feedback cycle, we replaced LacI with RNA-guided dCas9 (Fig. 1). With a suitable sgRNA, the dCas9-sgRNA complex represses TetR and thus fulfills the same functional role as LacI in the original circuit. We replaced LacI as this is the only endogenous protein from *E. coli* and might thus interfere with the circuit (previous repressilators were operated in the *lacI*-deficient strain MC4100). However, exchanging the P_lac_ promoter controlling the dCas9 gene with different constitutive promoters led to a deterioration or even disappearance of the oscillations (cf. SI Fig. S12). We therefore finally kept the P_lac_ promoter and either operated the oscillator in MC4100 or in MG1655 and supplementing the growth media with IPTG to alleviate the repression by endogenous LacI. We also found no GFP oscillation in a strain that lacks the protein degradation machinery. We thus left the degradation tags on *cI*-lambda and TetR and used strains with an active degradation machinery (*Materials and Methods*).

**FIG. 1.**
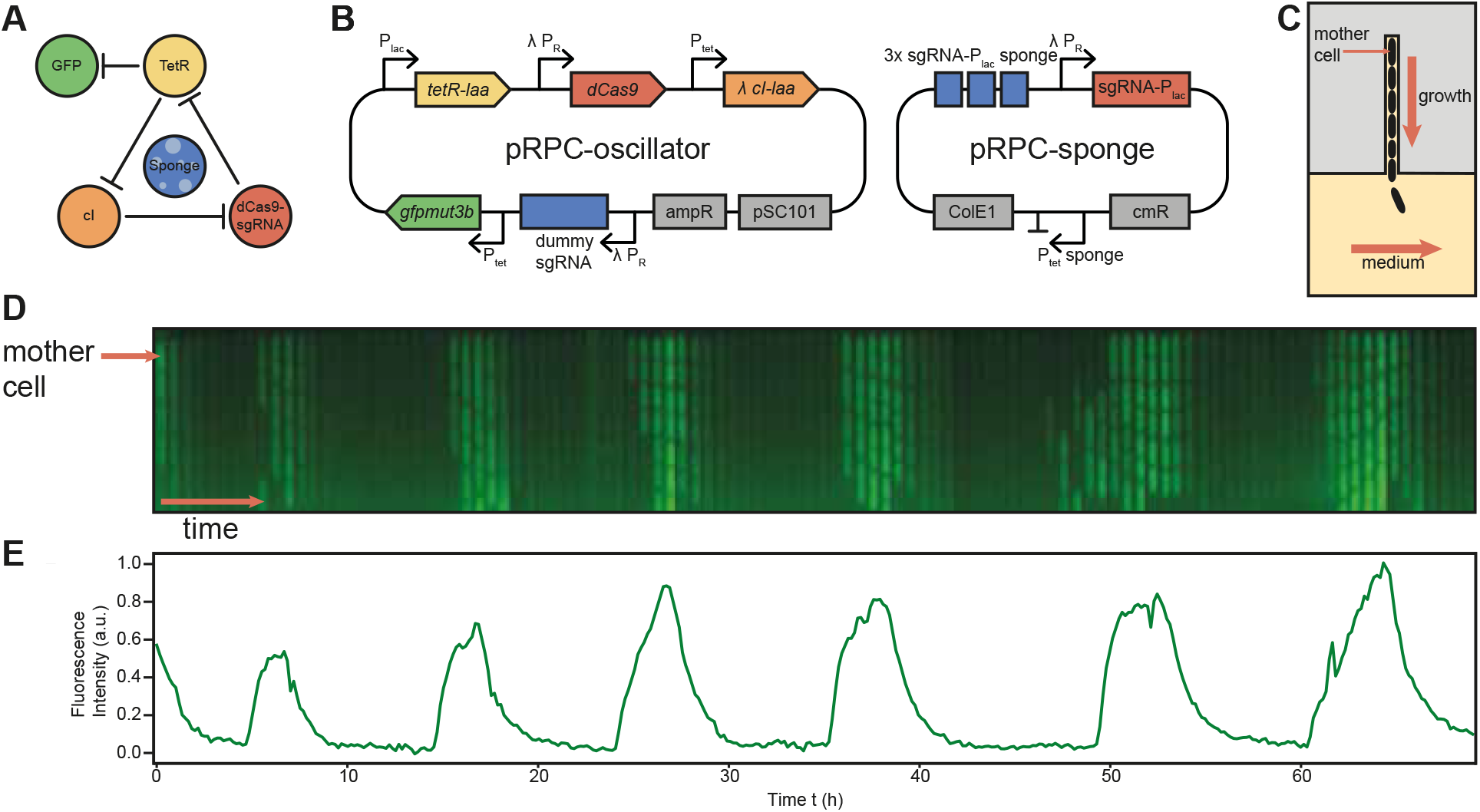
Design and microfluidic experiments. (A) Schematic representation of the repressilator circuit. A negative feedback loop of three repressors, including dCas9-sgRNA, oscillates, sponge elements stabilize the circuit dynamics and GFP is used as a fluorescent readout. (B) Schematic plasmid maps showing the realization of the genetic circuit on two plasmids with different copy numbers (pRPC-oscillator: ~5 (pSC101), pRPC-sponge: ~15-20 (ColE1). (C) Sketch of the mother machine microfluidic device. *E. coli* cells containing the plasmids are trapped in a small growth channel, where they are supplied with growth medium through a larger feed channel, allowing long-term observations of single cells in steady state growth conditions. (D) Kymograph (a single vertical growth channel over time) of cells displaying stable oscillations over almost 70 hours. (E) The fluorescence intensity (F.I.) of the corresponding mother cell over time is extracted from the microscopy videos by detecting the mother cell area and taking the average of the fluorescence intensity of that area.

Following the approach by Potvin-Trottier et al. [8], we introduced ‘sponge elements’ for the repressors, which were previously found to stabilize the dynamics against noise occurring at low copy numbers. To this end, we employed decoy binding sites for TetR and the dCas9-sgRNA complex on the sponge plasmid. We expressed an additional ‘dummy sgRNA’ from a *λ*-pR promoter on the oscillator plasmid. The latter acted as an additional sponge for *λ*-cI, while the dummy sgRNA competed with the formation of functional dCas9-sgRNA complexes (for experiments without sponge or dummy sgRNA cf. SI Figs. S10 & S11). The state of the oscillator can be tracked by a fluorescent protein (GFPmut3b) that is in phase with the *λ*-cI repressor.

### Single cell experiments

Since the oscillations are not actively synchronized across the bacterial population via the exchange of small molecule signals (such as in Ref. [24]), quantitative study of the oscillator requires the observation of the fluorescence intensity (F.I.) at the single cell level. Steady state growth conditions are also required for long term measurements, which can be achieved using a microfluidic growth chamber termed ‘mother machine’ [22] (Fig. 1C). In this microfluidic chip, a continuous supply of bacterial growth medium is pumped through a large channel (20 *μ*m × 50 *μ*m cross section). From this feed channel, small growth channels (1 *μ*m × 1 *μ*m cross section) branch off. The dimensions of the cross section allow only one *E. coli* cell to fit, effectively trapping single cells. Cell growth and division causes the growth channel to overflow into the feed channel, where the daughter cells are washed away. A single ‘mother cell’ at the end of the growth channel remains fixed in place and can be continuously observed under a microscope.

The data in Fig. 1 shows a typical example of a single growth channel filled with *E. coli* cells containing the two oscillator plasmids. A kymograph (Fig. 1D), displaying the fluorescence intensity of the selected growth channel over time, visualizes the periodic, oscillator-controlled GFP expression in the cells. The apparent synchronization of the cells is caused by their close relationship, as all cells within a single growth channel descend from the same mother cell and are thus expected to have similar cellular composition. The vertical position of the cells in the channel changes over time due to mechanical forces arising from the growth of the individual cells, which conversely also limits their growth rate [25]. Segmentation of the corresponding brightfield data (cf. Fig. S2) allows the extraction of quantitative fluorescence intensity data for the mother cell (Fig. 1 E). In this case, a total of six regular periods are visible over 69 hours of experiment.

### Statistical analysis of single cell data

Single cell analysis of a large number of individual mother cells from the same experiment allows the statistical analysis of the oscillator’s properties and contributes to a better functional understanding of its dynamics (Fig. 2). To this end, first a set of fluorescence intensity time traces {*I*_*i*_(*t*)} was extracted from the microscopy videos. All mother cells that showed normal growth in the bright field channel over the course of the experiment were included in the analysis. After 50 h of experiment run time, an increasing number of mother cells stopped dividing, hence the analysis is cut off after that time. In order to be able to better investigate the periodic portion of the noisy signal, we calculated the autocorrelation function (ACF) *A*_*i*_(*τ*) = 〈*I*_*i*_(*t* + *τ*)*I*_*i*_(*I*(*t*)〉_*t*_ for each mother cell. The ACF of the fluorescence intensity averaged over our set of single cell time traces, (i.e., 〈*A*〉(*τ*) = 1/*N* ∑_*i*_ *A*_*i*_(*τ*)) (Fig. 2A) shows that the analyzed cells oscillate on average. The first peak of the average ACF indicates an average oscillator period of 11.7 ± 0.4 hours (standard error of the mean, bootstrapped (SEM)). A histogram of individual oscillation periods (calculated from the peak-to-peak distances *T*_*j*_ for all full periods in {*I*_*i*_(*t*)}) also shows a mean of 11.7 h and a standard deviation of 3.5 h (Fig. 2B). The observed distribution has positive skewness (sample skewness g_1_ = 1.3), corresponding to the observation that most cells oscillate with a period in a relatively narrow interval around the mean (75% between 8h and 15h). Only a few oscillators were observed that ran very slowly, but no fast oscillations were recorded. Out of all mother cells that showed normal growth, only 5 out of 86 cells did not oscillate while for the remaining 81 cells 2 periods or more were registered. As shown in SI Fig. S1, within the same cell a given period *T*_*j*_ does not correlate with the previous period *T*_*j*−1_, indicating that the variance of the distribution is not exclusively explained by cell-to-cell variations, but also by fluctuations inherent to the synthetic circuit. We did not observe an overall slowing down of the oscillations, however, which conceivably could occur due to aging of the mother cell or other deterioration effects.

**FIG. 2.**
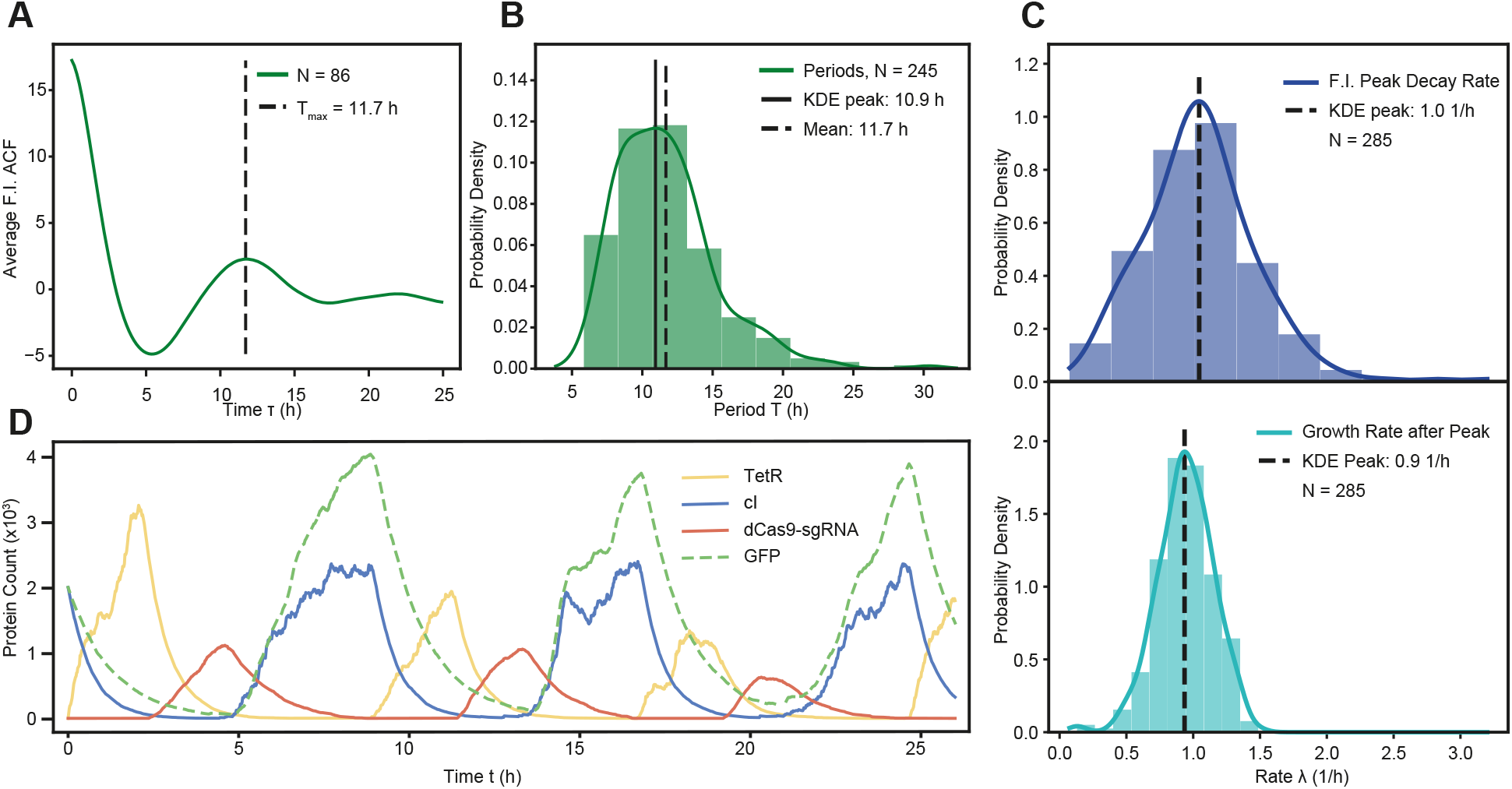
Analysis of single cell data. 86 channels measured in parallel over 50 h are analyzed and compared. (A) Autocorrelation function of the fluorescence intensity of single cells averaged over all mother cells. The first peak suggests that the time traces are periodic with a period of roughly 12 hours. (B) Distribution of periods (peak to peak distance) in all time traces (histogram with kernel density estimation (KDE)) shows the variation of the oscillator frequency. (C) F.I. peaks decay exponentially. The distribution of the rate (base e) over a fixed time interval is shown in on top. Comparison to the similar distribution of growth rates during the same interval suggests that dilution is responsible for protein decay. (D) Stochastic simulations match experimental data and reveal parameter constraints of oscillatory regime.

### Shape of the oscillations and relevant time-scales

The asymmetric shape of the peaks in the oscillator time traces suggests that our circuit behaves as a relaxation oscillator, which is consistent with previous findings for the repressilator v2.0 [8]. In a relaxation oscillator, a build-up phase of active protein production is followed by a decay phase, in which protein production has stopped and proteins are diluted or degraded. We find that the fluorescence time traces in the decay phases 1h – 4.5h after each peak are indeed closely matched by exponential fits (with a mean coefficient of determination 〈*R*^2^〉 = 0.94), as would be expected for relaxation oscillations.

In our oscillator circuit, the two repressor proteins are equipped with degradation tags, while the dCas9-sgRNA complexes are not actively degraded. We therefore expect dilution of dCas9-sgRNA (and potentially removal of the tightly bound complexes from their target sites by the replisome [26]) to be the slowest steps in the feedback circuit. Note that removal of dCas9-sgRNA would activate TetR, which in turn would suppress GFP production.

To elucidate the relation between the observed decay phase and dilution due to bacterial growth, we calculated an instantaneous growth rate *G*_*i*_(*t*) of the mother cell from division events observed in the microfluidic device (SI Fig. S4). From *G*_*i*_(*t*) we then obtained average growth rates for each decay period (i.e., 1h – 4.5h after each peak), and compared them to the observed fluorescence decay rates. Fig. 2 C shows histograms for both of these rates. The peaks decay with most likely rate *λ*_*dec*_ = 1.0 ± 0.03h^−1^ (SEM) while the cells grow with most likely rate of *λ*_*growth*_ = 0.9 ± 0.03h^−1^ (SEM). The decay of the fluorescence signal thus clearly occurs on the same time-scale as cell growth, but with a larger variability observed for the oscillator decay (CV = 40.4 %) than for the growth rate (CV = 22.3 %).

### Computer simulations of circuit dynamics

In order to check our assumptions about the dynamics of the RPC oscillator, we performed computer simulations using a stochastic model of the system. Using physically plausible parameters [27], the model results in time traces that closely recapitulate the experimental data (Fig. 2D). In particular the peculiar form of the oscillations and the time scale are matched very well. The model demonstrates that GFP production is out of phase with the presence of dCas9-sgRNA and roughly in phase with the production of *λ*-cI, as would be naively expected from the circuit topology. Accordingly, the period between peaks in GFP expression is determined by the decay phase of TetR, while the build-up phase of GFP is mainly determined by the dilution phase of dCas9-sgRNA.

It has been shown that dCas9-sgRNA binds to its target with immeasurably slow off-rates [28], and we therefore investigated the effect of such irreversible binding within our model. Our simulations suggest that irreversible binding of dCas9-sgRNA prolongs the dilution phase of dCas9-sRNA, which increases the period of the oscillations (cf. SI Fig. S13 & S14). The dilution of dCas9-sgRNA in turn is coupled to the replication and dilution dynamics of the circuit plasmids. Binding of *λ*-cI and LacI to their respective operators was assumed to be reversible (*Materials and Methods*).

In the simulation, the total number of sgRNA molecules, whose expression is controlled by *λ*-cI, limits the amount of dCas9-sgRNA formed. This keeps the build-up and subsequently also the decay of dCas9-sgRNA bounded in a parameter regime that allows for oscillations, which are, however, slower than in the original repressilator with LacI instead of dCas9-sgRNA [8]. The model is thus consistent with our assumption that removal of the dCas9-sgRNA complex is a rate-determining step of the oscillator. Furthermore, our model confirms that strong cooperativity is not required for the oscillations, but titration against the decoy binding sites on the sponge is important.

### Synchronization and loss of synchrony

Including a CRISPR node into a genetic oscillator suggests potential applications of such circuits for the control of secondary processes via CRISPRi that can be programmed by the choice of appropriate sgRNAs. In this context, the oscillator may either be used to periodically control the order of execution of the secondary processes (where keeping ‘absolute time’ may not be so crucial), or as an actual clock, which in combination with a molecular counter could be used to create precisely defined time delays. To evaluate the capability of our circuit to keep time, we experimentally measured how a set of initially synchronized single cell oscillators go out of phase over time and then used an information theoretic approach to interpret the data for the above applications.

Since we did not employ a mechanism for synchronization of oscillator cells via, e.g., quorum sensing-based communication [24], we initially synchronized the cells by stopping the oscillator using appropriate genetic inducers (Fig. 3A). Before the start of the single cell oscillator experiments, aTc was added to the medium used for bulk growth to stop repression by TetR, while IPTG was excluded from the medium, which would be normally required for the operation of the oscillator. After loading the cells into the mother machine devices, flow medium – now without aTc, but containing IPTG – was flushed in and used throughout the rest of the experiment. As all cells were thus locked in the same state at the beginning of the experiment, the oscillators were expected to start with the same oscillator phase.

**FIG. 3.**
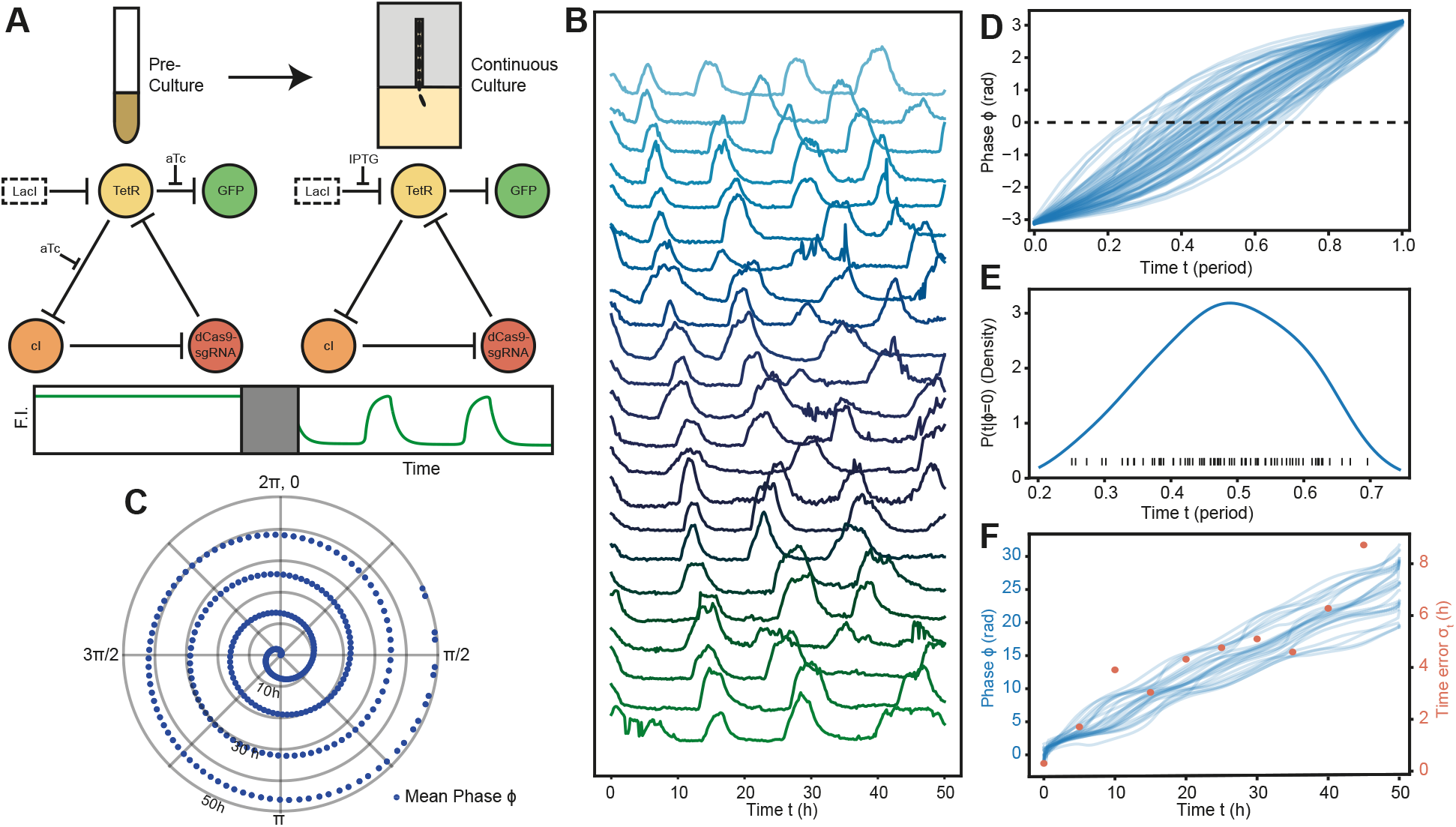
Dephasing behaviour of the oscillator. (A) Cells are synchronized by fixing the phase in the bulk pre-culture before the start of the microfluidic experiment. aTc is added to inhibit TetR and no IPTG is added to inhibit background of endogenous LacI, thus fixing the circuit in a state with high cI repressor and GFP concentration. After a time delay for experimental setup, the inducers are reversed in the microfluidic experiment the inducers are reversed and the cells begins oscillating. (B) Summary of single cell fluorescence intensity traces after phase synchronization. All traces start in phase at the tail end of a peak and dephase over time. (C) The instantaneous phase of the time traces is quantified with the Hilbert transform. The plot shows the average phase of all time traces on a circle with a radial time axis. (D) Variance of the instant. phase during a period is shown for all periods in the dataset, with time axis scaled by respective period length. Width of the conditional probability *P*(*t* |*ϕ* = 0) (inset, KDE and rug plot) determines how well different time states can be distinguished after measuring e.g. *ϕ* = 0 (w.l.o.g.). (E) Estimation of elapsed time after synchronization reduces in accuracy with loss of synchrony. Top shows set of instant. phase traces. Bottom shows a lower bound for the error in the time estimation at different time points.

Fig. 3B shows the fluorescence intensity time traces of all cells used in the analysis (cf. SI Fig. S7 for histograms). At the beginning of the traces, the cells appear to be in the decay phase after a (first) GFP maximum (this initial lag is explained by the time delay caused by transferring bacteria from culture to the mother machine and the start of timelapse acquisition (≈ 150 min)). To quantify dephasing among the oscillators, an instantaneous phase *ϕ*_*i*_(*t*) was calculated for every time point of every trace *I*_*i*_(*t*) by applying a Hilbert transform to the data (*Materials and Methods* and SI Fig. S3) [29]. From the distribution of the phase angles, we then obtained their mean and standard deviation at every time point (Fig. 3C & S6). The mean increases approximately linearly, with a slope corresponding to an average period length of 12.0 h. The standard deviation of the phase distribution steadily increases over time, and reaches half a period after 40 h, indicating a complete desynchronization of the observed set of oscillators in this time period.

### Reliability as a clock and pacemaker

Information-theoretic concepts have been previously applied successfully to gene regulatory as well as developmental processes [30, 31]. The fundamental question is whether the concentration of a biochemical species can be utilized by a biological system – in the presence of noise – as an estimator of some quantity of interest. In the case of gene regulation and sensing, the output of a genetic element would be used to ‘estimate’ the strength of a chemical or physical input, while in development it is used to infer spatial position. In analogy, we here ask, how reliable our synthetic clock could be used to tell time, or to control the order of molecular events dependent on the clock phase.

For the latter, a consistent temporal shape of the signal is required to allow a robust differentiation of distinct sections of an oscillation period. If one only cares about the order of events and not their absolute timing, the variation in the duration of the periods can be ignored. Hence we analyzed all full periods in the set of phase traces {*ϕ*_*i*_(*t*)}, normalized in time by their respective period length (Fig. 3D). To get a better idea of how to quantify the information the phase signal confers about time, let us consider the information gained by a single phase measurement. Before the measurement of the phase, the instant of time within one period is completely unknown (corresponding to a uniform prior distribution *P*(*t*)). After measuring a specific phase signal *ϕ*\*, the probability to be at a specific time point is then given by the conditional probability (i.e., posterior distribution) *P*(*t* |*ϕ*\*) (Fig. 3E). The narrower this distribution, the more uncertainty about the point in time is removed and hence more information is gained. This intuitive idea can be formalized using the concept of mutual information *I*(*t*; *ϕ*), which describes the amount of information about one variable (in bits) that is obtained from observing the other. From our experiments, we computed a value of 1.9 ± (5.8 · 10^−3^) bits (*Materials & Methods*), which indicates that theoretically a maximum of 2^1.9^ ≈ 3.7 distinct temporal states per period could be reliably distinguished from the output of a single node in an individual instantiation of the oscillator circuit.

For the application of the oscillator as a genuine ‘timekeeper’, it is interesting to ask how how precisely one could tell the absolute time passed since the initial synchronization of the bacteria (as in Fig. 3A,B) from the observation of an individual bacterial oscillator. As shown in Fig. 3F (blue), the variability in period length among the oscillators leads to a distribution of phases that spreads out over time. When the absolute time is estimated from the measurement of the phase, the variability in the phase signal, quantified by its standard deviation *σ*_*ϕ*_(*t*) leads to uncertainty..j of the time estimate *σ*_*t*_(*t*). A lower bound for *σ*_*t*_(*t*) is given by the Cramer-Rao bound 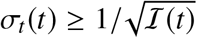, where 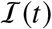 is the Fisher information calculated from the phase angle distribution (*Materials & Methods*). As the oscillators go out of phase, the uncertainty about the absolute point in time increases (Fig. 3F, red), i.e., the accuracy of the clock decreases. While the absolute error increases almost four-fold over the course of the experiment (compare *σ*_*t*_(*t* = 5*h*) ≥ 1.7*h* with *σ*_*t*_(*t* = 40*h*) ≥ 6.5*h*), the error relative to the runtime of the experiment actually goes down.

We need to point out that the above considerations somewhat artificially refer to the phase data mathematically extracted from the fluorescence signal generated by the oscillator. In order to actually ‘read out’ the oscillator phase, the bacteria would have to make use of instantaneous protein levels and also their rates of change. Further, our analysis of the phase shown in Fig. 3D assumes that the bacteria ‘know’ when an oscillation period starts or ends (corresponding to a peak in protein expression), which makes the phase uncertainty vanish at these points. Deriving temporal information from several of the oscillator nodes (rather than from just one), might further improve the ability to determine a point in time within an oscillation. Finally, in order to be able to measure *absolute* time (as a timekeeper), the oscillator circuit would have to be coupled to a counter [32] (otherwise the accumulated phase *ϕ*(*t*) is not accessible, but only *ϕ*(*t*) mod 2*π*).

### Interplay with growth dynamics

Gene expression activity is intimately linked to the bacterial growth rate [33, 34], and it can therefore be expected that genetic oscillations will be correlated with periodic changes in cell growth. Indeed, our single cell analysis revealed that the instantaneous growth rate *G*_*i*_(*t*) of each mother cell oscillates in phase with and with the same frequency as the fluorescence intensity. Strong periodic fluctuations in the cell growth are already visible in the single cell growth data (cf. Fig. S9 for example traces). To deal with the relatively noisy growth data, we computed the cross-correlation *C*_*i*_(*τ*) = 〈*G*_*i*_(*t* + *τ*)*I*_*i*_(*I*(*t*)〉_*t*_ between the growth and fluorescence traces for each single cell and then averaged over the whole set of cells. *C*_*i*_(*τ*) measures the similarity of the two signals as a function of the relative time shift *τ* between them. Ideally, the cross-correlation of two synchronous signals peaks at *τ* = 0 and at the common period *τ* = *T*. While the ACF of the fluorescence intensity for our synchronized cells oscillates with a period of 12.4 h (cf. ACF in Fig. S8), the average cross-correlation data in Fig. 4 peaks at *τ* = 0 h and *τ* = 12.7 h, demonstrating that the instantaneous growth rate indeed oscillates in phase with the fluorescence intensity. We point out that bacterial cell division occurs on a different time-scale than these oscillations – the average instantaneous growth rate in the experiments was 1.13 ± 0.44 h^−1^, corresponding to a doubling time of 53 min with a *CV* = 39% (see SI Fig. S8). We also performed control experiments with bacteria, in which the oscillations were permanently suppressed as shown in Fig. 3A. In this case, no correlation was obtained between growth rate and fluorescence (Fig. 4), further supporting that the experimentally observed dynamic changes in growth rate are indeed caused by the oscillations of the circuit.

**FIG. 4.**
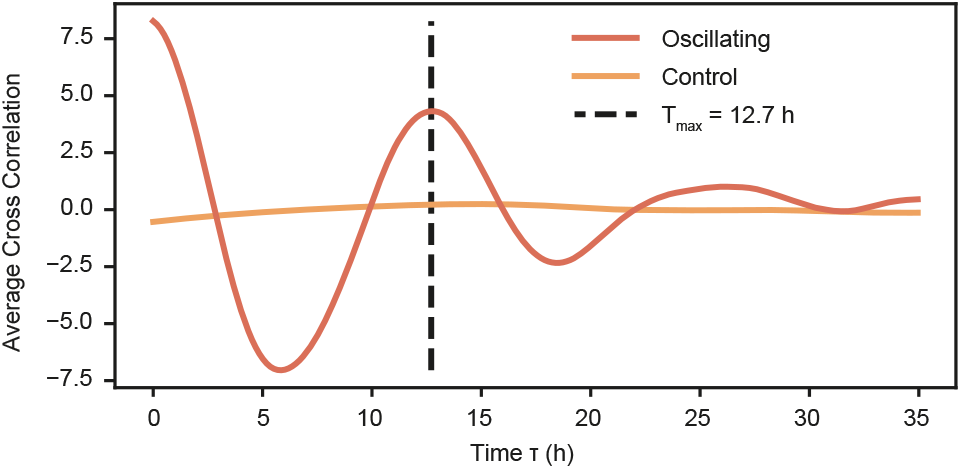
The cross-correlation of fluorescence intensity *I*(*t*) and instantaneous growth rate *G*(*t*) averaged over the set of mother cells shows the growth rate oscillates in phase with the fluorescence intensity. A control experiment with limited oscillations (aTc added, no IPTG, see Fig. 3A) shows no correlation between fluorescence intensity and instantantaneous growth rate.

There are manifold ways in which oscillator dynamics and growth could be coupled. In our oscillator circuit, the production of GFP is in phase with the production of the *λ*-cI repressor, which represses the expression of sgRNA and dCas9. Since high GFP fluorescence is correlated with a high instantaneous growth rate, an influence of dCas9-sgRNA expression and impeded growth appears likely [35, 36]. It is further conceivable, that interactions between the circuit and bacterial growth feed back on the oscillator dynamics in a complex way: a reduction in growth speed would also reduce the dilution rate of the oscillator components, which in turn would affect the timing of the relaxation oscillations.

## Discussion and Conclusion

In this work, we have integrated transcriptional repression via CRISPR interference into a three-node repressilator network (termed the ‘RPC oscillator’), which resulted in stable genetic oscillations at the single cell level. A statistical analysis of a large number of bacterial oscillators showed considerable variability in the periods among the cells, but also fluctuations of the periods within individual cells. The shape of the oscillatory fluorescence readout signal indicates that the RPC oscillator functions as a relaxation oscillator. Stochastic numerical simulations suggest that titration of the oscillator components against decoy binding sites as well as the dilution or removal of dCas9-sgRNA complexes from their target sites due to cell growth play a decisive role in its dynamics. This also explains why – with a period of ≈ 12h – the RPC oscillator is relatively slow compared to other previously published synthetic bacterial oscillators. Quite notably, we found that the genetic oscillations of the RPC oscillator were accompanied by periodic changes of the cellular growth rate with the same period. We surmise that this effect might be caused by the periodic cellular load generated by dCas9-gRNA expression during operation of oscillator, indicating that the use of CRISPRi in dynamical circuits requires special care.

On the other hand, the sequence-programmable knockdown of gene expression via CRISPRi opens up a wide range of potential applications, as secondary processes controlled by this synthetic clock need not be constrained to gene expression from plasmids. It is easily conceivable that the CRISPRi-coupled oscillators could be used to control endogenous, chromosomal genes by simply supplying the corresponding sgRNAs.

In order to explore the potential of the RPC circuit to act as a temporal controller or timekeeper, we analyzed the cell-to-cell variability of the oscillator phase within an information-theoretic framework. The results suggest that the output of a single oscillator node could be used to reliably distinguish three to four phases within a single oscillator period, which could be used, e.g., to control the order of the corresponding number of downstream processes. The oscillator could also be used as a real ‘clock’ to measure ‘absolute time’ starting from a defined point in time. We found that due to fluctuations in the oscillator phase, an initially synchronized population of oscillators loses synchrony over time – after about 40 hours (or four oscillation periods) the error in time is on the order of 6 hours (or half a period).

Based on the above considerations, future studies will aim at further improving the dynamics of the RPC oscillator and explore potential applications. The load on the cells could be reduced by utilizing a genome-integrated gene for dCas9, which potentially would reduce the observed influence on cellular growth. An interesting opportunity could be the use of sgRNA-aptamer fusions [37], which would make the CRISPRi process dependent on the presence of the aptamer’s target molecule. This could be potentially used to tune the oscillator frequency, and also to reset or entrain the synthetic clock.

Rather than using repression by CRISPRi, one could also use CRISPR activation (CRISPRa) [38] based on dCas9-activator fusions, both as part of the oscillator circuit or for the control of secondary processes. The most exciting prospect would be to control bacterial gene expression of chromosomal genes with the oscillator, which could be achieved by targeting these genes with the corresponding sgRNAs. This could be utilized, e.g., to couple the oscillator to metabolic processes such as the utilization of certain metabolites. By supplying these metabolites in a rhythmic manner, this in turn could be used to evolve ‘circadian’ oscillators with a desired frequency.

## Supplementary Information (SI)

The SI contains Supplementary Methods, SI Figs. S1-S14, SI Tables S1-S3, SI References.

## MATERIALS AND METHODS

### Cloning

The two plasmids used in this publication were constructed using standard PCR, restriction ligation and Gibson assembly [39]. DNA sequences are listed in the SI. The pRPC-oscillator plasmid is based on pZS1-lTlrLLtCL [3] (Addgene #26489) with the dCas9 gene taken from pdCas9-bacteria [18] (Addgene #44249). The pRPC-sponge plasmid is based on pLPT41 [8]. Cloning steps were performed in *DH5α*.

### Lithography

A master mold for the ‘mother machine’ microfluidic device was fabricated in a cleanroom by lithography. Four layers of SU8-2000 negative photoresist (Microchem) with varying thickness (corresponding to the desired channel height) were applied and exposed consecutively on a 2 inch silicon wafer (Siegert Wafer): Firstly, a base layer (roughly 1 μm height) helps the adhesion of fine structures in subsequent layers. A second layer with alignment markers is used as position reference for the subsequent two layers. The growth channels (20-30 μm length) are patterned in thin third layer (1 μm height). Feed channels (150 μm width) are patterned in a fourth layer (20 μm height). In the case of the third layer, the resist was crosslinked using electron-beam lithography (Raith eLINE, 30 kV acceleration voltage, 10 μm aperture). The other three layers were UV-crosslinked in a mask aligner (Süss MicroTec, 200W lamp) with photomasks (Zitzmann). The lithography process parameters (dilutions, spin, bake, development and exposure parameters) that were found to produce these specifications are given in a detailed protocol in the SI.

### Bacterial cell culture

Bacterial cultures (K-12 wild type strain *MG1655* containing the plasmids described above) were grown overnight in LB medium, diluted 1 to 100 into 10 ml LB medium and grown for another 3 hours (37 °C, 250 rpm). For loading of the microfluidic device, the cultures were spun down (5000 rcf for 5 min) and resuspended in 150 μl flow medium (LB medium supplemented with 1.1 ‰ (w/v) bovine serum albumin (BSA) for passivation). All medium was supplemented with carbenicillin (50 μg/ml) and chloramphenicol (12.5 μg/ml) antibiotics. If specified, the inducers isopropyl *β*-D-1-thiogalactopyranoside (IPTG) and anhydrotetracycline (aTc) were added (500 μM and 214 nM, respectively).

### Microfluidics

To construct the microfluidic devices, PDMS and curing agent (Dow Corning, Sylgard 184) were mixed (1:10) and poured on the master wafer in an aluminium foil container, degassed for 30 min and baked for 80 min at 80 °C. The cured PDMS was detached from the master wafer and individual chips were cut out. The channel side of the chip was immediately covered with Scotch magic tape to keep it clean. Inlet and outlet holes were punched with a biopsy puncher. Cover slips (170 *μ* m thickness, Carl Roth, LH26.1) were cleaned with Hellmanex III (2 %, Hellma) and rinsed with ddH_2_O. PDMS chips were bonded to the cover slips after O_2_ plasma treatment (0.7 min, Diener Femto) and baked for 60 min at 80°C to complete the bonding. The complete microfluidic devices were loaded with highly concentrated bacterial culture with a syringe through a PTFE tube (0.8 mm inner diameter, Bola, S1810-10) attached to the outlet. The loading was observed under a microscope and the pressure on the syringe adjusted so that the flow of the culture stopped. The bacteria were left to settle for 30 min (at 37 °C). After that the loading syringe was cut off, the free end of the tube connected to the outlet was placed into a tube for waste collection and the dense loading culture was flushed out with flow medium from a 20 ml syringe through a second silicone tube attached to the inlet. A continuous flow of flow medium was set up with a syringe pump (0.5 ml/h, TSE Systems) for the duration of the time lapse. Syringes were replaced as needed taking care not to introduce air bubbles. In the case that a buildup of bacteria stuck in the feed channel was observed in the feed channel during a timelapse, a manual flush was performed periodically.

### Microscopy

Time lapse videos were recorded with a Nikon Ti-2E microscope using a 60x plan apochromat oil objective (NA 1.40), 1.5x magnification, a SOLA SM II LED light source, Andor NEO 5.5 camera and NIS elements software. The temperature in the microscope enclosure was controlled at 37 °C, the microfluidic chip and tubing were fixed in place with custom 3D printed clamps for optimal stability. The Nikon Perfect Focus System was used to account for focus drift. For the timelapse, up to 40 different positions were imaged in phase contrast brightfield and GFP fluorescence channels for 200 ms every 10 minutes for up to 70 hours.

### Data Analysis

Timelapse videos were split per imaged position and cropped to selected growth channels with custom imageJ [40] macros (selection criterium: successful loading and continuous growth observed the brightfield channel over the course of the timelapse). A segmentation mask for trapped cells was created with the software ilastik [41] (pixel classification using a random forest classifier, all input features selected). Training of the classifier was done by specifying the class (cell or background) of sample pixels. To achieve good results, examples of all relevant image features were included in the training, while taking care not to overtrain the classifier on specific examples. All following data analysis steps were done with Python (Version 3.7, Python Software Foundation) using the numpy, scipy and pandas libraries. Segmentation masks were cleaned up with binary morphological operations, connected pixels corresponding to the mother cell were determined by position in the channel, fluorescence intensity values were averaged over mother cell pixels and corrected with reference to the should-be constant value of PDMS background. In cases where the mother cell stopped dividing during the timelapse, the first daughter cell below the mother cell was used as substitute. The instantaneous growth rate was calculated by detecting cell division events and differentiating smoothed (Savitzky-Golay filter) cumulative division event data. To calculate the instantaneous phase, the phase angle of the analytical representation of the fluorescence intensity data was calculated and unwrapped. This (complex) analytical signal was constructed with the smoothed fluorescence intensity signal as real part and the Hilbert transform thereof as imaginary part [29, 42]:

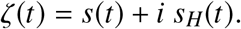

The Hilbert transform of a real signal is defined by:

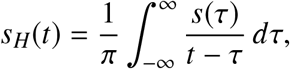

where the integral is understood as its Cauchy principal value. The analytical signal has a positive spectrum with no negative frequency component, and it can be represented by complex vectors rotating in one direction. The instantaneous phase is then readily available as the angle of the analytical signal: *ϕ*(*t*) = arg [*ζ*(*t*)]

The concept of mutual information is based on Shannon’s information entropy [43]:

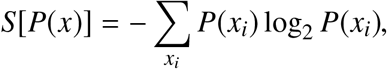

which measures the uncertainty about a random variable *x* ∈ {*x*_*i*_} with probability distribution *P*(*x*_*i*_). The mutual information quantifies the statistical dependence between two random variables and their associated probability distributions and thus the amount of information that can be gained about one variable by observing the other, which can be expressed as:

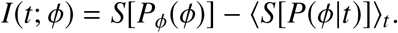

Written in this way, it can be interpreted as the information contained in the dynamic range of the measured signal minus the average information entropy of the variability of the signal at fixed time points, i.e., noise. To estimate a value for the mutual information from our experimental data, we used the *second Gaussian approximation* described by Tkačik et al. [31]. Briefly, the conditional probability distributions *P*(*ϕ*|*t*) were approximated as Gaussian and the marginal distribution *P*_*ϕ*_(*ϕ*) is obtained by integrating the conditional prob. dist. over time. The effect of the a limited number of samples is dealt with by extrapolating to infinite samples from randomly sampled subsets of the available data (cf. Fig. S5).

The Cramer-Rao bound [44] gives a local value about the error of the time estimate from knowledge of the phase (compare to the global value of the mutual information). It states:

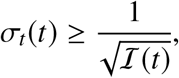

with the Fisher information 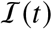 (for a Gaussian *P*(*ϕ*|*t*)) given by

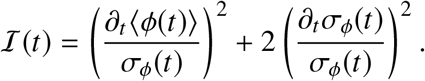

The phase averaged over the set of cells 〈*ϕ*(*t*)〉 and the corresponding standard deviation *σ*_*ϕ*_ was calculated directly from the phase data (see above); the derivatives were smoothed (Savitzky-Golay filter).

The standard error of the mean for values that correspond to peak locations (ACF/histogram) was estimated from a bootstrapped sample distribution (random resampling of the data with replacement, 1000 times).

### Simulation

Stochastic simulations were based on a model by Potvin-Trottier *et al.* [8]. The simulation approximates the reactions in one cell based on realistic parameter values that have been found previously (see SI). The simulation does not consider changes in the cell volume due to growth and division, nor the partitioning of molecules at cell division. In order to match experimental and simulated GFP traces, we optimized parameters and quantities such as the concentrations of TetR, *λ*-cI and dCas9-sgRNA. The simulations were performed using a Gillespie algorithm [45], with propensities for the expression of *λ*-cI, dCas9, sgRNA, dummy sgRNA and GFP given by

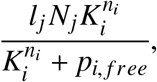

where *l*_*j*_ is the expression rate of molecule *j* that is repressed by molecule *i*, *N*_*j*_ is the number genes for *j*, *K*_*i*_ is the repression threshold of molecule *i*, *n*_*i*_ the hill coefficient and *p*_*i, free*_ is the amount of unbound molecules *i*. In case of irreversible binding of dCas9-sgRNA to DNA target sites the propensity for TetR expression is given by *l* · *N*_*free*_ where *l* is the expression rate and *N*_*free*_ is the amount of genes without bound dCas9-sgRNA. Otherwise the propensity for TetR expression was calculated with the Hill function as described above (see SI. Proteins are produced in bursts of size *b* = 10, whereas transcription of sgRNA results in one new molecule. sgRNA and dCas9 expression is simulated separately followed by a reaction to form the active molecule dCas9-sgRNA. All proteins, including dCas9-sgRNA, are degraded by dilution according to the growth rate. Due to their degradation tags, TetR and *λ*-cI are additionally degraded enzymatically. sgRNA is exclusively degraded by enzymes at a time scale much shorter than dilution. The maturation of GFP was included as a separate reaction.

## ACKNOWLEDGMENTS

We gratefully acknowledge financial support by the European Research Council (grant agreement no. 694410, project AEDNA) and the BMBF through the EraSynbio network (project UNACS, grant no. 031L0011). M.S.-S. acknowledges support by the DFG through the GRK 2062. The pLPT41 plasmid was a kind gift from Laurent Potvin-Trottier and Johan Paulsson.

