## Supporting Information for "Single cell characterization of a synthetic bacterial clock with a hybrid feedback loop containing dCas9-sgRNA"

**CONTENTS**

Supporting Methods

Figures S1-S14

Tables S1-S3

Supporting references

### SUPPORTING METHODS

#### Master Fabrication Protocol

- 4 layer lithography
- SU-8 2000 resist
- 2 inch wafer

Base Layer:

- Do not clean wafer
- Spin coat 3ml SU-8 2000.5
  - 500/100/10 (rpm speed/(rpm/sec) acceleration/sec duration)
  - 3000/300/60
  - Observe regular circular interference pattern, otherwise discard wafer
- Soft bake 1 min @ 95 °C (hotplate)
- UV exposure 15 sec (200 W mask aligner, soft contact) with no photo mask
- Post exposure bake 1 min @ 95 °C

Alignment Marker:

- Spin coat 3ml SU-8 2002
  - 500/100/10
  - 3000/300/60
- Soft bake 3 min @ 95 °C
- UV exposure 15 sec 30 sec (200 W mask aligner, soft contact) with alignment marker photo mask
- Post exposure bake 3 min @ 95 °C
- Develop for 30 sec, gentle agitation in SU-8 developer

- Rinse wafer for 10 sec with fresh SU-8 developer, 10 sec with isopropyl alcohol. Dry with pressured air.

Growth Channels:

- Spin coat 3ml SU-8 2000.5
  - 500/100/10
  - 3000/300/60
- Soft bake
  - 1 min @ 65 °C (65 °C steps optional)
  - 3 min @ 95 °C
  - 1 min @ 65 °C
- e-beam exposure with eLINE, take care about the following:
  - loadlock procedure
  - dose (exposure screen necessary)
  - alignment and rotation
  - focus (use scratch on wafer), stigmation, aperture parameters
  - writefield alignment
  - structure design
- Post exposure bake
  - 1 min @ 65 °C
  - 3 min @ 95 °C
  - 1 min @ 65 °C
- Develop for 30 sec, gentle agitation in SU-8 developer
- Rinse wafer for 10 sec with fresh SU-8 developer, 10 sec with isopropyl alcohol. Dry with pressured air.

- Hard bake 15 min @ 150 °C optional

Feed Channels:

- Spin coat 3ml SU-8 2025
  - 500/100/10
  - 5000/300/60
- Soft bake
  - 1 min @ 65 °C
  - 4 min @ 95 °C
  - 1 min @ 65 °C
- Locally develop alignment markers by cleaning resist with swab soaked with SU-8 developer. Mark the locations of alignment markers before spin coating last layer if alignment markers are not visible.
- UV exposure 45 sec (200 W mask aligner, hard contact) with feed channel photo mask aligned to the alignment markers on the wafer. Alignment critical for correct growth channel length.
- Post exposure bake
  - 1 min @ 65 °C
  - 4 min @ 95 °C
  - 1 min @ 65 °C
- Develop for 90 sec, gentle agitation in SU-8 developer
- Rinse wafer for 10 sec with fresh SU-8 developer, 10 sec with isopropyl alcohol. Dry with pressured air.
- Hard bake 15 min @ 150 °C

### SUPPORTING TABLES AND FIGURES

TABLE I. Overview over microfluidic experiments used in this publication. Experiments listed in the table all use the plasmids described in the main text in strain *MG1655*. Kymographs in SI Figs. show exeriments with varying plasmid versions and strains.

| Dataset | Label | Loaded Channels | BF Select. | FI Select. | $t_{\text{cutoff}}$ | Fig. |
|---|---|---|---|---|---|---|
| 1 | Main | 383 | 86 | 86 | 50 h | 2 |
| 2 | Sync. | 129 | 60 | 23 | 50 h (70 h) | 1, 3, 4 |
| 3 | Control | - | 54 | 54 | 35 h | 4 |
| 4 | Prelim. | 73 | 1 | 1 | | |

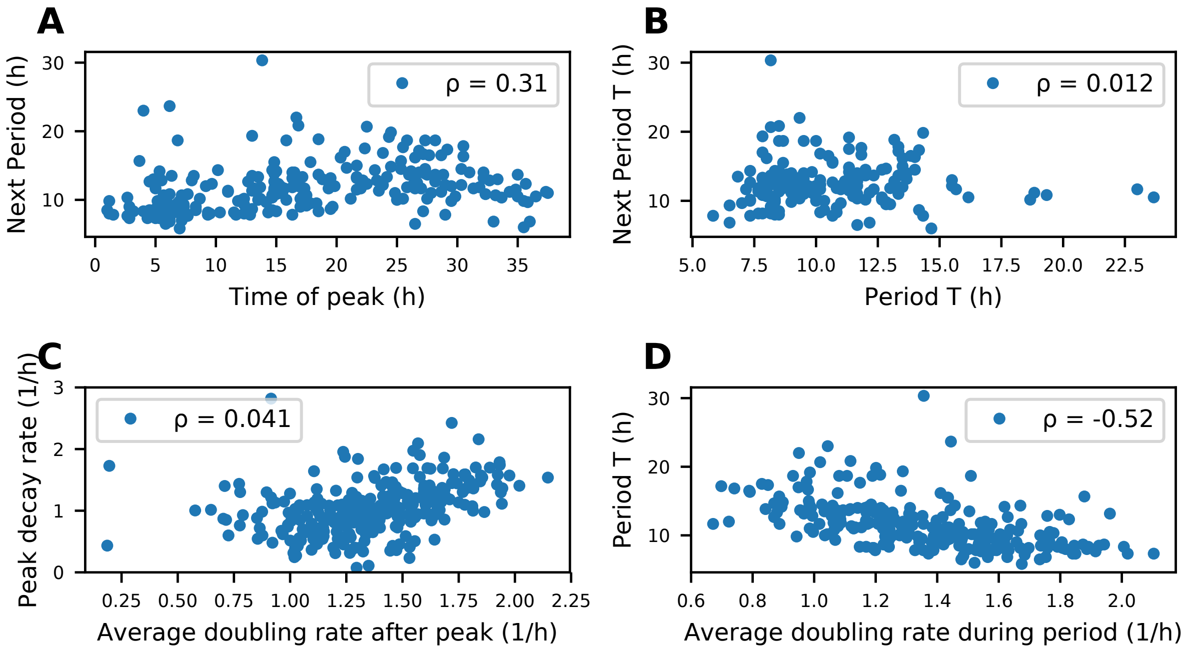


FIG. 1. Scatter plots showing the relationships between different quantities calculated from the single cell data (dataset 1). Pearson correlation coefficient $\rho$ is given in the legend. (A) Time of a F.I. peak against the duration of the following period. A slowdown of the oscillator is not observed. (B) Length of a period against the length of the following period in the same mother cell. (C) Average doubling rate after a peak (1h - 4.5h) against the peak decay rate in the same interval. (D) Average doubling rate during an entire period against the length of the period.

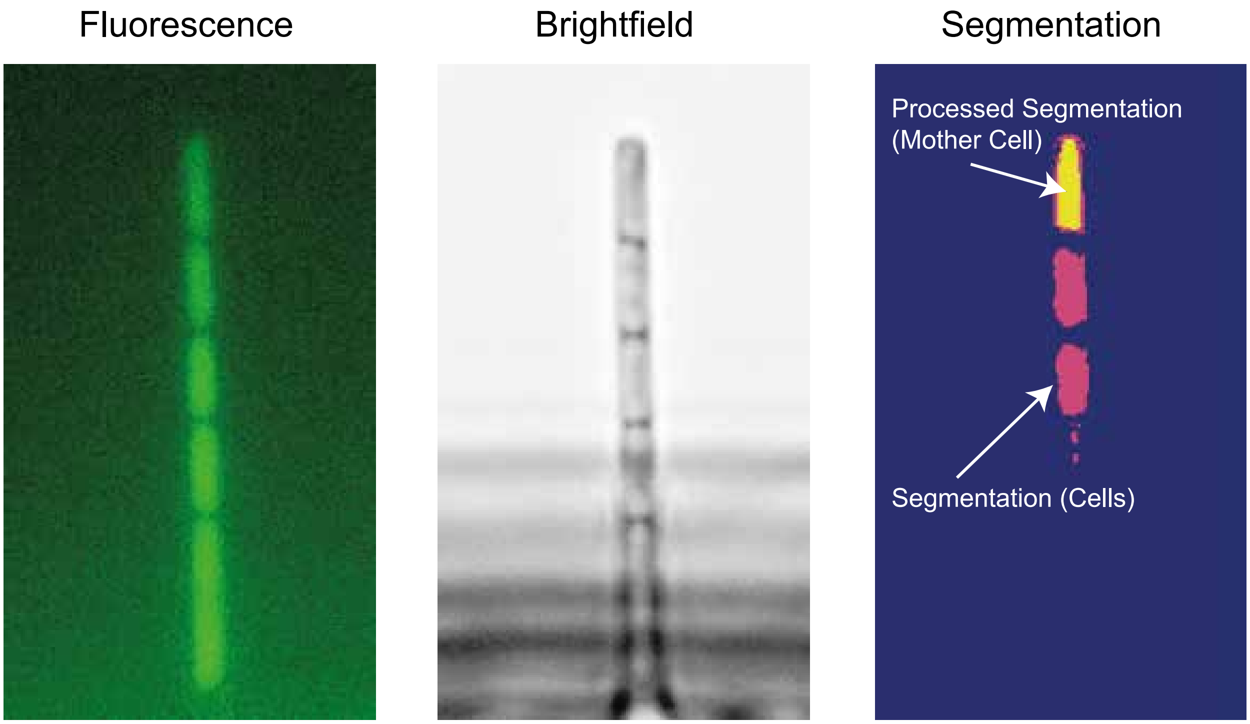


FIG. 2. Sketch of the segmentation process used to extract single cell data from microscopy videos. The brightfield channel (B) is used to create a segmentation mask corresponding to cells using ilastik (cells in the lower part of the growth channel are ignored). The initial segmentation is processed to refine the mask and detect the mother cell. The resulting mother cell binary mask is applied to the fluorescence intensity channel and values for the average over the cell area are extracted for each frame.

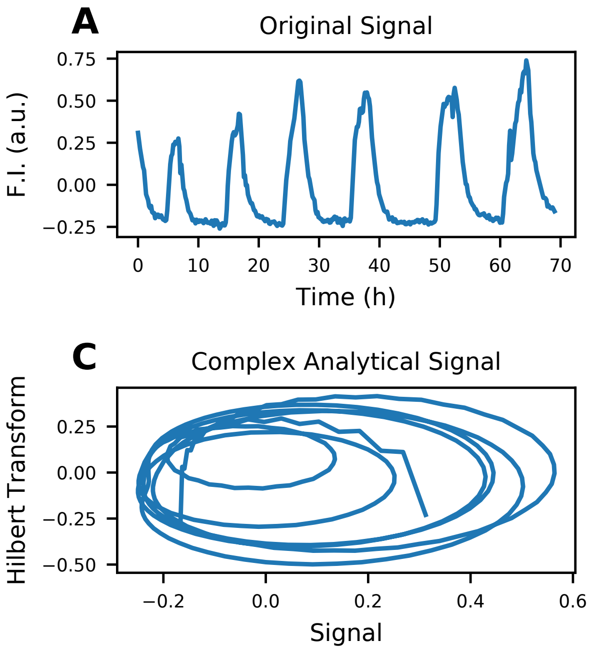


FIG. 3. Illustration of the calculation of the instantaneous phase. The oscillating signal (A) is smoothed (B) with a Savitzky-Golay filter. The Hilbert transform is used to construct a complex analytical signal. A visualisation in the complex plane (C) shows the circular trace of the analytical signal vector. The instantaneous phase is the angle of the complex vector (D). The phase angle ( mod $2\pi$) can be unwrapped to measure the progression over multiple periods.

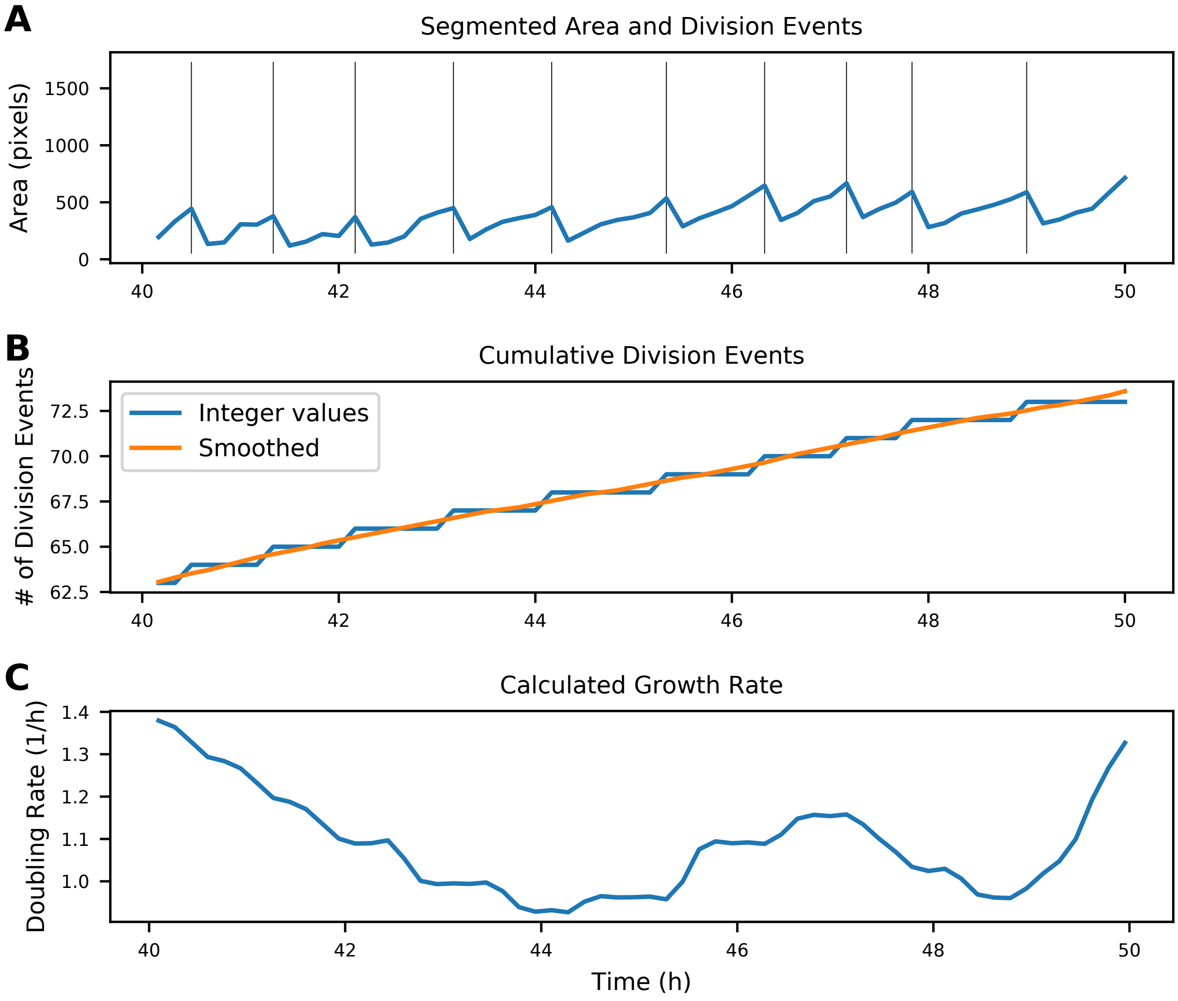


FIG. 4. Illustration of the calculation of the instantaneous growth rate. (A) The area of the segmentation mask for the mother cell is used to detect division events (vertical line). (B) The discrete division events are added up to generate a curve of cumulative division events. This curved is smoothed to effectively interpolate between the discrete division events. (C) The derivative of the smoothed curve is the instantaneous growth rate (doubling rate).

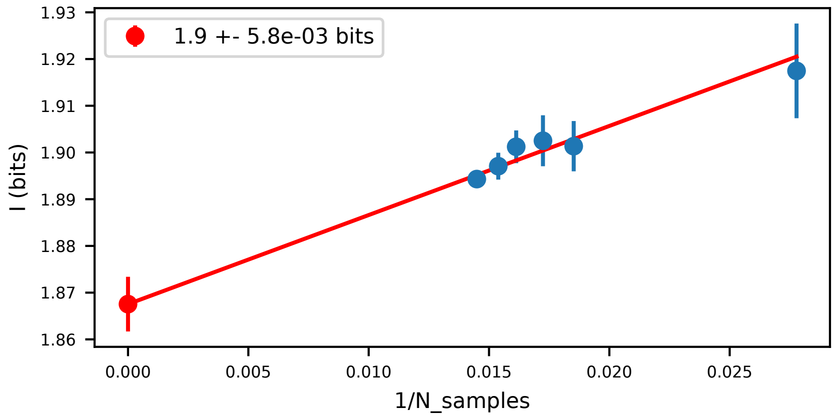


FIG. 5. Extrapolation of the value for the mutual information (Fig. 3D) to an infinite number of samples. The mutual information is calculated for subsets of the original data of different sizes (random resampling without replacement). Each size of subset is resampled 100 times to calculate mean and standard error. The y-intercept of a linear fit corresponds to an extrapolated value for an infinite number of samples.

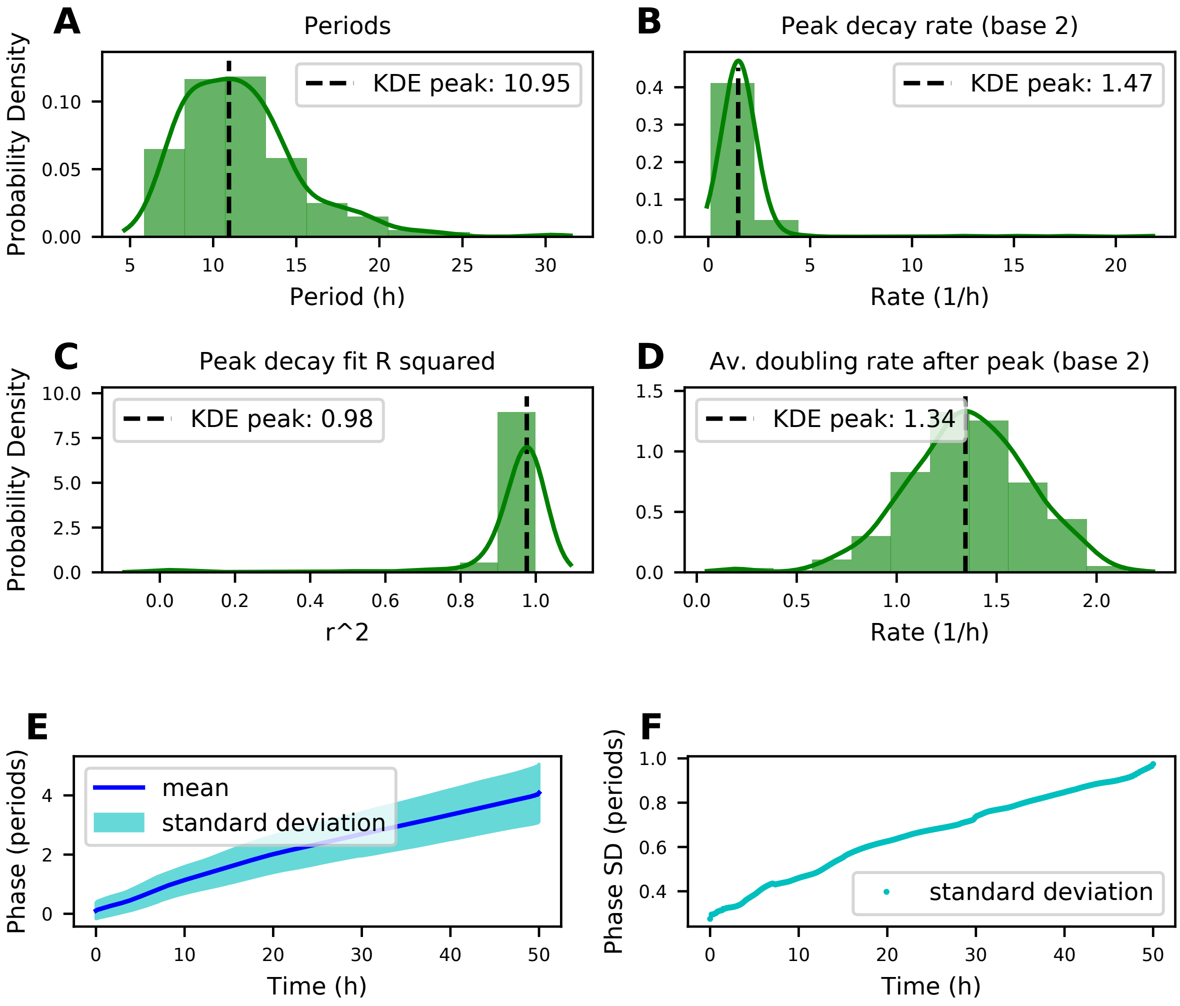


FIG. 6. Dataset 1: Overview. (A) Histogram of periods in the dataset. (B) Histogram of the decay rate of the peaks (1h to 4.5h after a peak). (C) Histogram of the coefficient of determination of the fits in (B). (D) Average doubling rate 1h to 4.5h after a peak. (E) Instantaneous phase over time, average (with standard deviation) over all cells in the dataset. (D) Standard deviation of (E) over time.

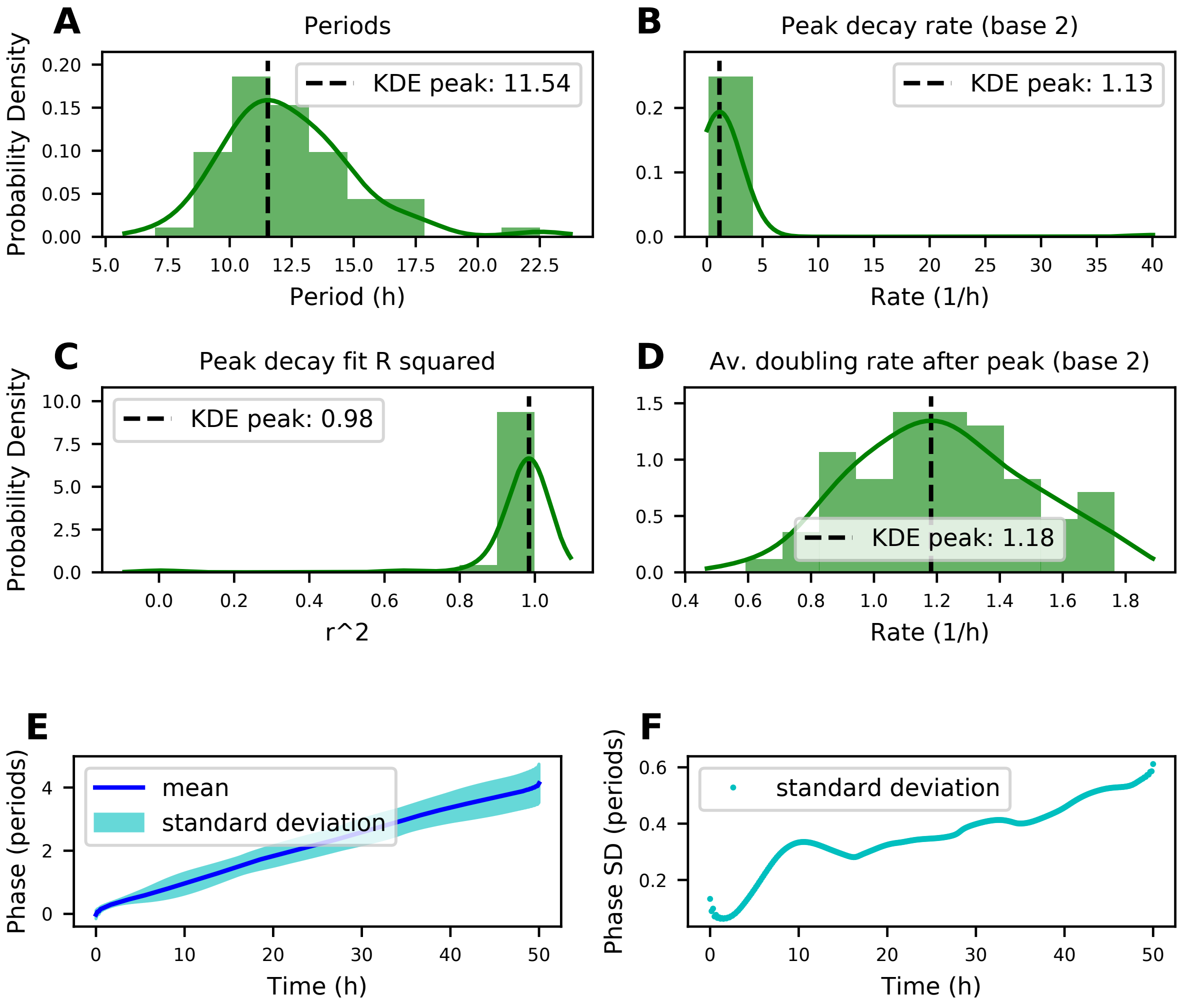


FIG. 7. Dataset 2: Overview. (A) Histogram of periods in the dataset. (B) Histogram of the decay rate of the peaks (1h to 4.5h after a peak). (C) Histogram of the coefficient of determination of the fits in (B). (D) Average doubling rate 1h to 4.5h after a peak. (E) Instantaneous phase over time, average (with standard deviation) over all cells in the dataset. (D) Standard deviation of (E) over time.

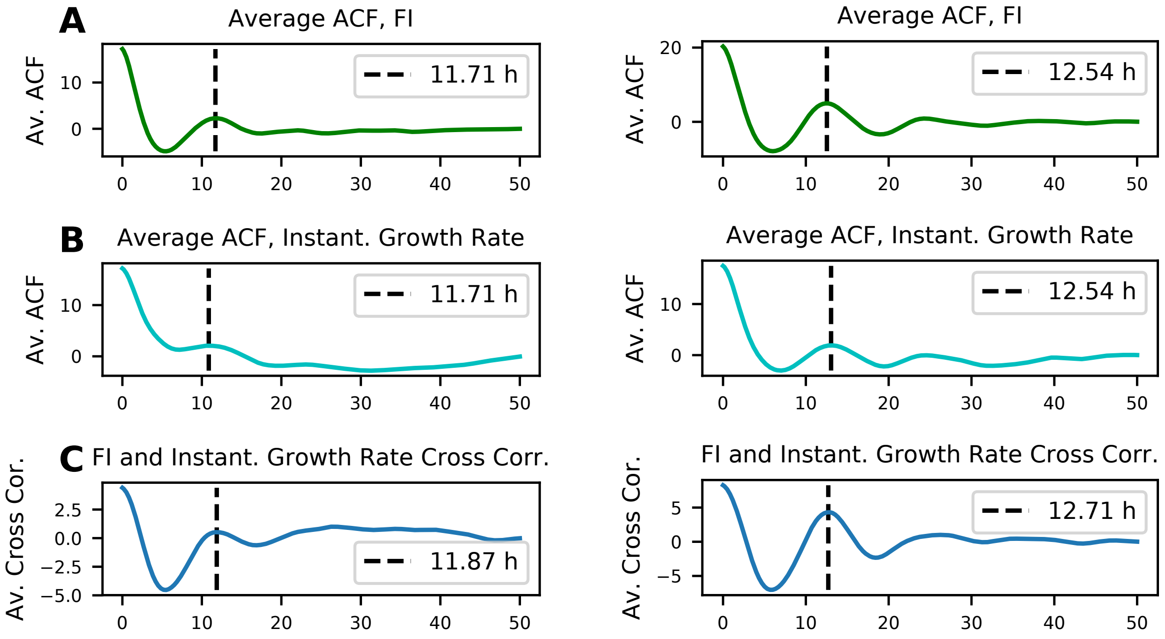


FIG. 8. Correlation functions for dataset 1 (left) and dataset 2 (right). (A) Average autocorrelation function of the fluorescence intensity. (B) Average autocorrelation function of the instantanous growth rate reveals oscillations similar to F.I. (C) Average cross correlation between instant. growth rate and F.I. confirms both signals oscillate in phase.

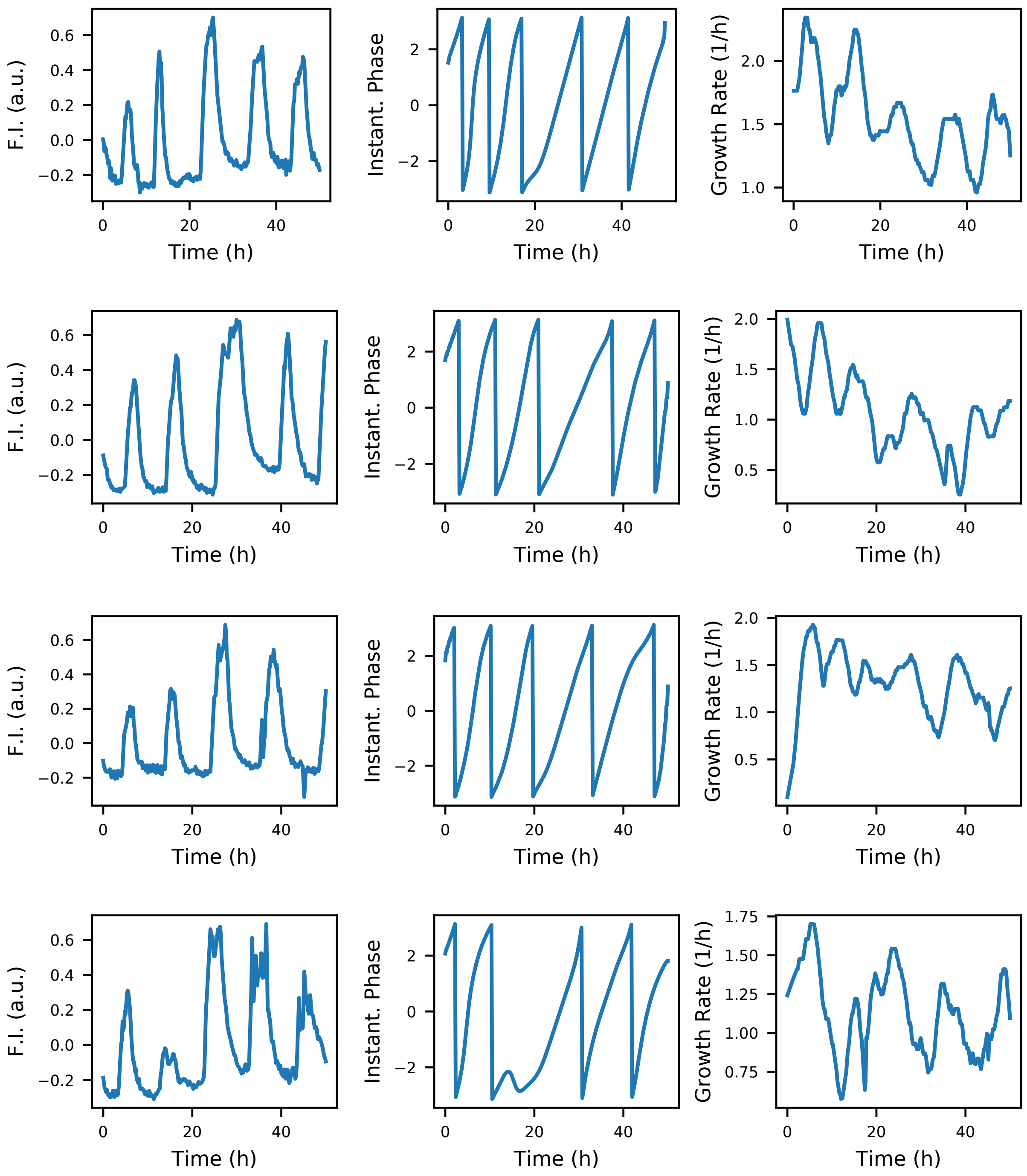


FIG. 9. Dataset 1 example time traces for GFP fluorescence intensity (left), instantaneous phase (wrapped, middle) and growth (doubling) rate (right).

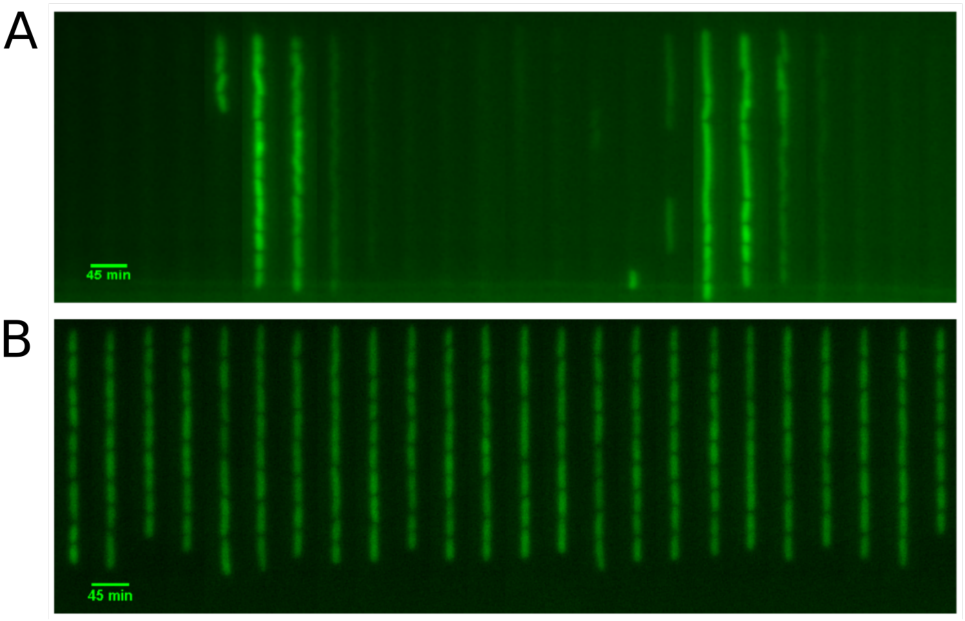


FIG. 10. **A** Kymograph (a single vertical growth channel over time - supply channel at the bottom) of cells displaying stable oscillations with a version of the RPC plasmid that does not contain the dummy sgRNA. The sponge plasmid is the same except that it lacks the sponge element for the dummy sgRNA. **B** Kymograph (a single vertical growth channel over time - supply channel at the bottom) of cells with the RPC plasmid without the dummy sgRNA and without a sponge plasmid. The control experiment shows that without the sgRNA for the dCas9 to target TetR which is encoded on the sponge plasmid there are no oscillations. However, from 44 analyzed cells about 43% still show a detectable GFP signal. In both experiments the *E. coli* strain MC4100 was used since it lacks LacI.

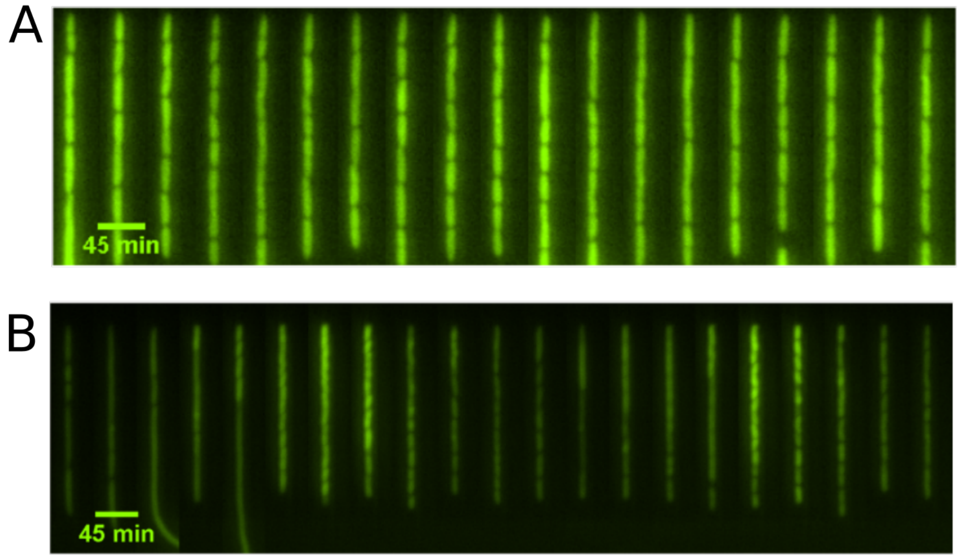


FIG. 11. **A** Kymograph (a single vertical growth channel over time - supply channel at the bottom) of cells displaying stable oscillations with a version of the RPC plasmid that does not contain the dummy sgRNA. Furthermore we exchanged the $P_L lacO1$ with the constitutive promoter pJ23101. The bacteria stopped oscillating. **B** Kymograph (a single vertical growth channel over time - supply channel at the bottom) of cells with the RPC plasmid that does not contain the dummy sgRNA. Furthermore we exchanged the $P_L lacO1$ with the constitutive promoter pJ23106. The oscillations of the GFP concentration in the cells reduced in intensity and are less regular. In both experiments the *E. coli* strain MC4100 was used which lacks LacI. The sequences for the promoters were taken from the Berkeley 2006 iGEM team (parts.igem.org/Part:BBa_J23101).

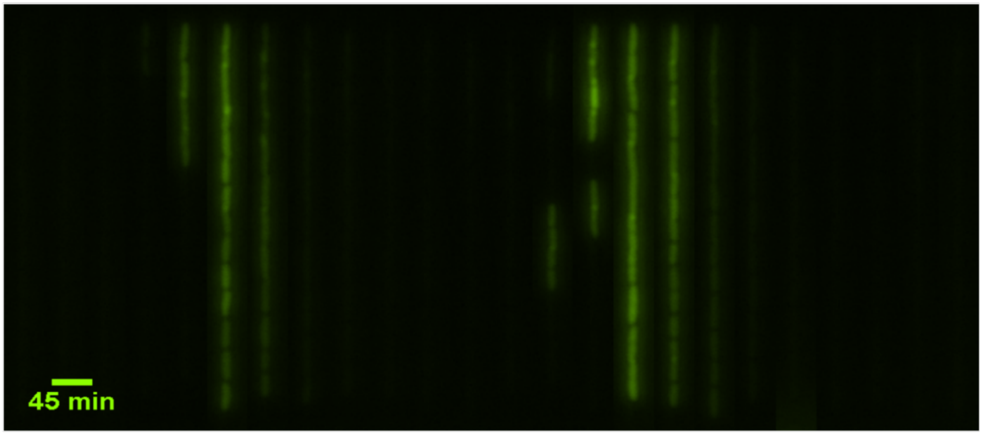


FIG. 12. Kymograph (a single vertical growth channel over time - supply channel at the bottom) of the *E. coli* strain MC4100 displaying stable oscillations with the RPC plasmid with the dummy sgRNA and the sponge plasmid. The *E. coli* strain MC4100 was used which lacks LacI.

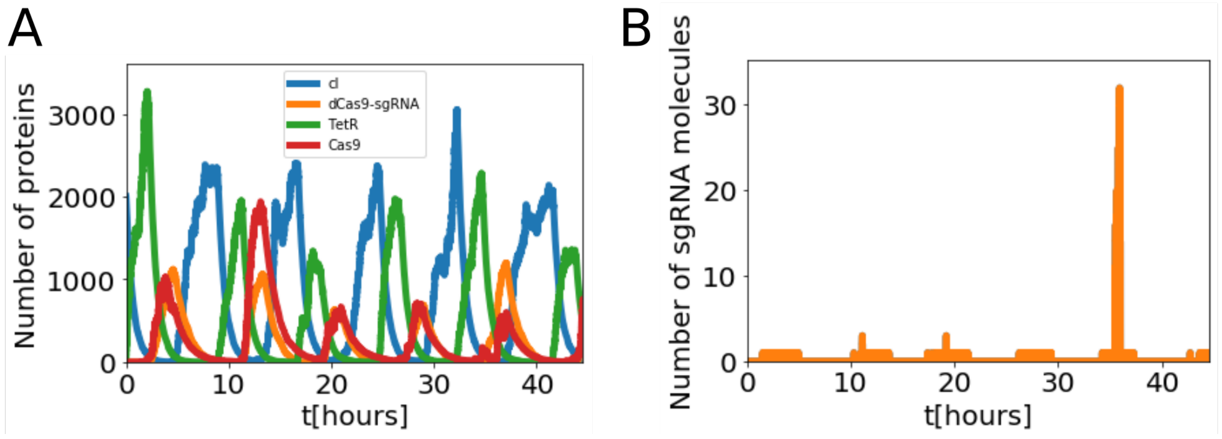


FIG. 13. **A** Simulated time traces of number of proteins from the genetic oscillator. The amount of dCas9-sgRNA complex is not limited by the amount of dCas9. **B** Simulated time trace of the number of sgRNA molecules for the same simulation run as in (A). sgRNA molecules limit the number of dCas9-sgRNA molecules that form. The simulation assumes irreversible binidng of dCas9-sgRNA to its target sites.

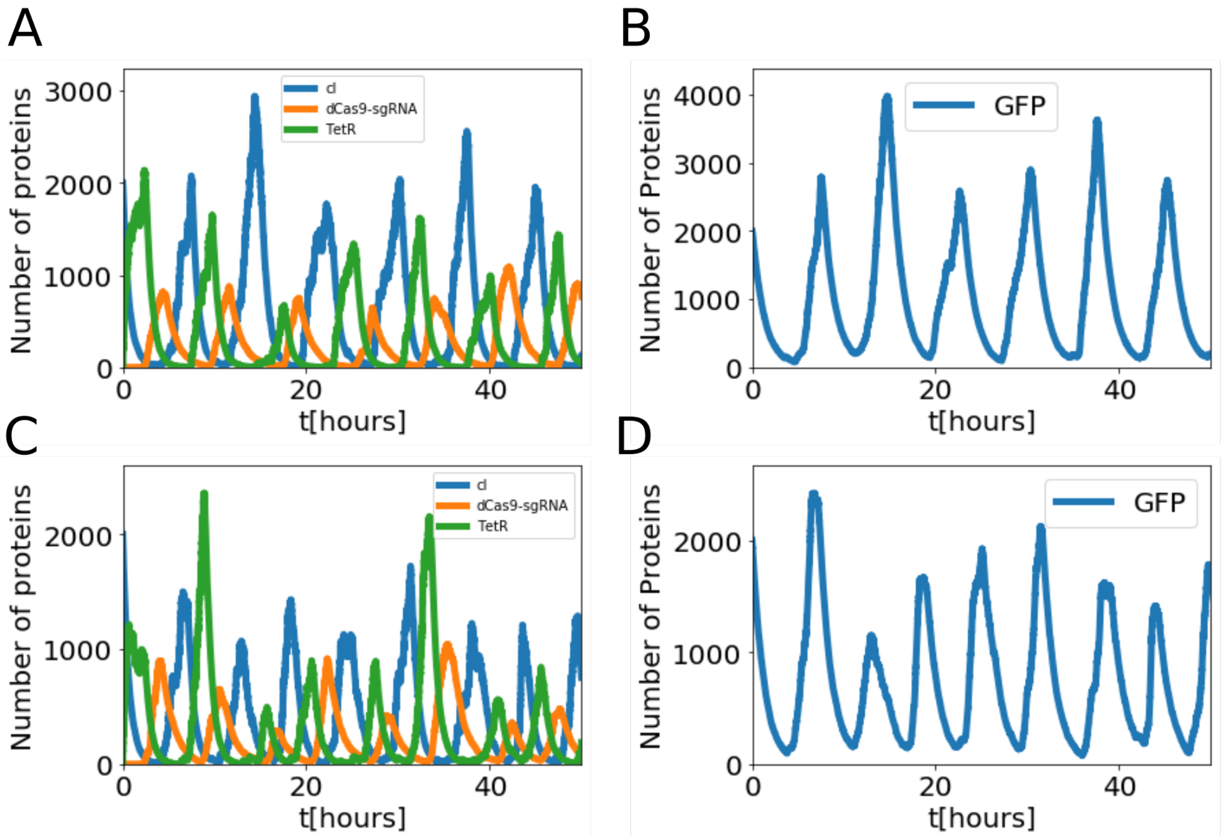


FIG. 14. Simulation of the oscillator, where the repression of all three proteins including dCas9-sgRNA is simulated by a Hill function. **A** Simulated time traces of number of proteins from the genetic oscillator with a threshold value of $K = 0.1nM$ and a hill coefficient of $n = 1$ for dCas9-sgRNA. **B** Simulated time traces of number of GFP molecules from the genetic oscillator for the simulation run as in (A).**C** Simulated time traces of number of proteins from the genetic oscillator with a threshold value of $K = 1nM$ and a hill coefficient of $n = 1$ for dCas9-sgRNA. **D** Simulated time traces of number of GFP molecules from the genetic oscillator for the simulation run as in (C).

**STOCHASTIC MODEL**

### List of reactions

Our stochastic simulation of the oscillator using a Gillespie algorithm is based on the model from Potvin-Trottier *et al.* [1]. The list of reaction with their corresponding propensities are shown below. Here we assume irreversible binding of dCas9-sgRNA to the DNA target sites.

$\mathrm{i} \xrightarrow{l_i \cdot (N_{rep} + N_{rep+dCas9}) \cdot} \mathrm{K}_j^{n_j}\ \mathrm{K}_j^{n_j} + \mathrm{j}_{free}\ \ \mathrm{i} + \mathrm{b}$, for $i = (dCas9, cI)$ and $j = (cI, TetR)$

$\mathrm{TetR} \xrightarrow{l_{TetR} \cdot N_{rep}} \mathrm{TetR} + \mathrm{b}$

$\mathrm{GFP} \xrightarrow{l_{GFP} \cdot (N_{rep} + N_{rep+dCas9}) \cdot} \mathrm{K}_{TetR}^{n_{TetR}}\ \mathrm{K}_{TetR}^{n_{TetR}} + \mathrm{TetR}_{free}\ \ \mathrm{GFP} + \mathrm{b}$

$\mathrm{sgRNA} \xrightarrow{l_{sgRNA} \cdot (N_{sponge} + N_{sponge+dCas9(1,2,3)}) \cdot} \mathrm{K}_{cI}^{n_{cI}}\ \mathrm{K}_{cI}^{n_{cI}} + \mathrm{cI}_{free}\ \ \mathrm{sgRNA} + 1$, with $N_{sponge+dCas9(1,2,3)} = N_{sponge+1 \cdot dCas9} + N_{sponge+2 \cdot dCas9} + N_{sponge+3 \cdot dCas9}$

$\mathrm{sgRNA}_{\mathrm{dummy}} \xrightarrow{l_{sgRNA} \cdot (N_{rep} + N_{rep+dCas9}) \cdot} \mathrm{K}_{cI}^{n_{cI}}\ \mathrm{K}_{cI}^{n_{cI}} + \mathrm{cI}_{free}\ \ \mathrm{sgRNA}_{\mathrm{dummy}} + 1$

$\mathrm{dCas9} + \mathrm{sgRNA}_{\mathrm{i}} \xrightarrow{l_{complex} \cdot i \cdot dCas9} \mathrm{dCas9{-}sgRNA}_{\mathrm{i}}$, with $i = (sgRNA, sgRNA_{dummy})$

$\mathrm{i} \xrightarrow{l_{dil} \cdot i} \mathrm{i} - 1$, with $i = (\mathrm{dCas9}, \mathrm{dCas9} - sgRNA, \mathrm{dCas9} - sgRNA_{dummy}, GFP, GFPmat, N_{rep} + N_{rep+dCas9}, N_{sponge} + N_{sponge+dCas9(1,2,3)})$

$\mathrm{i} \xrightarrow{(l_{dil} + l_{tag}) \cdot i} \mathrm{i} - 1$, with $i = (cI, TetR)$

$\mathrm{i} \xrightarrow{l_{deg} \cdot i} \mathrm{i} - 1$, with $i = (sgRNA, sgRNA_{dummy})$

$\mathrm{i} \xrightarrow{l_{dil} \cdot < N_i >} \mathrm{i} + 1$, with $i = (N_{rep}, N_{sponge})$

$\mathrm{dCas9{-}sgRNA} + \mathrm{N_i} \xrightarrow{l_{target} \cdot \mathrm{dCas9} - sgRNA \cdot N_i} \mathrm{dCas9{-}sgRNA{-}1, N_i{-}1 and N_j} + 1$,
with $i = (N_{rep}, N_{sponge}, N_{sponge+1 \cdot dCas9}, N_{sponge+2 \cdot dCas9})$

and $j = (N_{rep+dCas9}, N_{sponge+1\cdot dCas9}, N_{sponge+2\cdot dCas9}, N_{sponge+3\cdot dCas9})$

GFP $\xrightarrow{l_{mat}\cdot GFP}$ GFPmat + 1

$p_{free}$ is calculated by solving the following equation: $p_{tot} = p_{free} + N \cdot \frac{p_{free}^n}{p_{free}^n + K^n}$ where $N$ is the total number of binding site on the DNA and $p$ the corresponding repressor protein.

$< N_i >$ is the copy number of the corresponding plasmids.

TABLE II. Simulation parameters.

| **Rate constant** | **Value** | **Source** |
|---|---|---|
| Transcription rate | $l_{sgRNA} = 0.02s^{-1}$ | [2] |
| RNA degradation | $l_{deg} = 0.01s^{-1}$ | [2] |
| Gene expression rate | $l = 0.0189s^{-1}$ | [1] |
| Protein dilution rate due to growth | $l_{dil} = ln(2)/(53 * 60)s^{-1}$ | Measurement |
| Protein degradation due to tag | $l_{tag} = 0.0002s^{-1}$ | [3] |
| dCas9-sgRNA complex formation rate | $l_{dCas9-sgRNA} = 6.11 \cdot 10^6 s^{-1}$ | [4] |
| Translation bursts | $b = 10$ | [1] |
| GFP maturation time | $l_{mat} = 3.33 \cdot 10^{-3} s^{-1}$ | [5] |
| Hill coefficients for cI, TetR | $n = 1.9$ and $n = 1.2$ | [6] |
| Repression constants for cI, TetR | $K = 5.9nM$ and $K = 2.3nM$ | [6] |
| Repression constant and hill coefficient for dCas9-sgRNA | $K = 0.1 - 1nM$ and $n = 1$ | Estimation |
| Binding rate of dCas9-sgRNA to DNA | $l_{target} = 1.191 \cdot 10^7 s^{-1}$ and $n = 1$ | [4] |
| Copy number repressilator | $N_{repressilator} = 5$ | [7] |
| Copy number sponge | $N_{sponge} = 20$ | [8] |

| Parent | Template | Assembly | Product | Sequence Name | Sequence |
|---|---|---|---|---|---|
| pZS1-lTlrLLtCL | BBa I13522 | AatII, XhoI | pRep-GFP | I13522_fwd | tgccacctgacgtctaagaa |
| | | | | I13522_rev | gcggacgctcgagattaccgcctttgagtgagc |
| pRep-GFP | pdCas9-bacteria | Gibson | pRPC-base | pdCas9_fwd | ggttgcatgtactagcgacagatctaaagaggagaaaggatctatgg |
| | | | | pdCas9_rev | gagctgtcttcggtatcctaggtataaacgcagaaaggc |
| | pRep-GFP | | | pRep-GFP_fwd | atacctaggataccgaagacagctcatg |
| | | | | pRep-GFP_rev | agatctgtcgctagtacatgcaaccattatcacc |
| pRPC-base | gBlock | AatII | pRPC-oscillator | sgRNAd_insert | tgccacctgacgtctaagaagtcgaggataaatatctaacaccgtgcgtgttgactattttacctctggcggtgataatggttgcccgtccagctcgaccagaatgttttagagctagaaatagcaagttaaaataaggctagtccgttatcaacttgaaaaagtggcaccgagtcggtgctttttttccaggcatcaaataaaacgaaaggctcagtcgaaagactgggcctttcgttttatctgttgtttgtcggtgaacgctctctactagagtcacactggctcaccttcgggtgggcctttctgcgtttatagacgtcgctcactcaaaggcggtaat |
| pRPC-oscillator | gBlock | ClaI, SbfI | pRPC-J23106 | dLacI_J23106_ins | gcaatccatcgattttacggctagctcagtcctaggtatagtgctagctcaaagacattccaaccacgttacgattaaagaggagaaaggtaccatgtccagattagataaaagtaaagtgattaacagcgcattagagctgcttaatgaggtcggaatcgaaggtttaacaacccgtaaactcgcccagaagctaggtgtagagcagcctacattgtattggcatgtaaaaaataagcgggctttgctcgacgccttagccattgagatgttagataggcaccatactcacttttgccctttagaaggggaaagctggcaagattttttacgtaataacgctaaaagttttagatgtgctttactaagtcatcgcgatggagcaaaagtacatttaggtacacggcctacagaaaaacagtatgaaactctcgaaaatcaattagcctttttatgccaacaaggtttttcactagagaatgcattatatgcactcagcgctgtggggcattttactttaggttgcgtattggaagatcaagagcatcaagtcgctaaagaagaaagggaaacacctactactgatagtatgccgccattattacgacaagctatcgaattatttgatcaccaaggtgcagagccagccttcttattcggccttgaattgatcatatgcggattagaaaaacaacttaaatgtgaaagtgggtctgcagcaaacgacgaaaactacgctttagcagcttaatctagaggcatcaaataaaacgaaaggctcagtcgaaagactgggcctttcgttttatctgttgtttgtcggtgaacgctctcctgagtaggacaaatccgccgccctagacctagctgcaggtcgaggataaatatctaacaccgtgcgtgcctgcagggcaatcc |
| pRPC-J23106 | pRPC-J23106 | Overhang PCR | pRPC-J23101 | dJ23106_J23101_fwd | ctaggtattatgctagcaagacattccaaccacgttac |
| | | | | dJ23106_J23101_rev | gactgagctagctgtaaaagctgataccgctcgcc |
| pLPT41 | gBlock | NotI, SacI | pLPT41-sgRNA | sgRNA-expression_insert | atgcttaaatcgcggccgctcaataactcaattgagagcatcaagctgcaggtcgaggataaatatctaacaccgtgcgtgttgactattttacctctggcggtgataatggttgcatctaacatctcaatggctagttttagagctagaaatagcaagttaaaataaggctagtccgttatcaacttgaaaaagtggcaccgagtcggtgctttttttccaggcatcaaataaaacgaaaggctcagtcgaaagactgggcctttcgttttatctgttgtttgtcggtgaacgctctctactagagtcacactggctcaccttcgggtgggcctttctgcgtttatagtgaacgagctcgcatatctcg |
| pLPT-sgRNA | gBlock | SpeI, NotI | pRPC-kan-sponge | sgRNA-sponge_insert | ctcaattgagaactagtgcatcaaccttagccattgagatgttagatgtatggaagcaattgcgcgggtgagagggtaattagccatcttcaccaacataacgctatcgggtcatactataagattccacttagtgaatgcaactatttataccttagccattgagatgttagattattctaagttacacttagtgtaatcatttttcatcagatttgctatagcccttgaacactacatgcatgaaaccaaattatgtatacactgggtcattaataccttagccattgagatgttagattcgcctacaattacctacaacggtcgaccatgcataccttcgattatcgtggccactctcggattacagcggccgccggcagaagtg |
| pRPC-kan-sponge | pSB1C3 | SacI, AatII | pRPC-sponge | | |

Table S3: Overview of DNA sequences used in the construction of plasmids used in this work.
